# Bolero: a dedicated workflow to decipher Hepatitis B virus transcriptome from long-reads sequencing method coupled to 5’RACE amplification of transcripts

**DOI:** 10.1101/2024.09.17.613398

**Authors:** Xavier Grand, Guillaume Giraud, Alia Rifki, Alexia Paturel, Delphine Bousquet, Armando Andres Roca Suarez, Cyril Bourgeois, Massimo Levrero, Fabien Zoulim, Barbara Testoni

## Abstract

Hepatitis B virus (HBV) represents a major health burden, as it affects close to 290 million people worldwide. Although prophylactic vaccines are available, current therapeutic compounds do not usually achieve HBV eradication due to the persistence of the covalently closed circular (ccc)DNA that serves as viral reservoir. Thus, novel biomarkers that reliably reflect intrahepatic cccDNA transcriptional activity would be highly relevant for the monitoring of infected individuals, as well as the evaluation of new treatments targeting HBV. In this context, the development of 5’ rapid amplification of complementary DNA ends (5’RACE) as a strategy to capture and amplify full-length HBV RNAs, coupled with long-read and full-length sequencing approaches (e.g., Oxford Nanopore Technology), has recently enabled the detailed characterization of these molecules. The analysis of such data requires a dedicated bioinformatics pipeline due to the highly condensed nature of the HBV genome, which is characterized by the production of multiple transcripts and spliced variants that overlap each other. Here, we present Bolero, a computational method and built-in workflow designed to handle HBV sequencing data and evaluate the relative expression of viral RNAs and their spliced variants. The analysis of HBV-infected cell lines demonstrates that our bioinformatics pipeline is efficient for the identification and quantification of individual HBV mRNAs. Thus, Bolero represents a useful tool to study cccDNA transcriptional activity and the heterogeneity of HBV RNA spliced variants.

**Author summary:** Transcriptomic analyses have brought comprehensive insights in the mechanisms controlling gene expression. Moreover, with the recent advances in sequencing technologies and computational methods, researchers can nowadays not only quantify gene expression, but also study alternative splicing, polyadenylation, transcription initiation, and even rare phenomena such as distant gene fusions. However, conventional analysis tools still rely heavily on the assumption of linear genomes with minimal overlap between open reading frames, rendering them insufficient for studying complex viruses such as hepatitis B virus (HBV).

Unlike typical linear genomes, HBV genome consists in a circular DNA molecule, which results in an extensive sequence overlap between its transcripts. To tackle these challenges, we developed an innovative approach coupling 5’ rapid amplification of complementary DNA ends (5’RACE) and long-read sequencing to comprehensively explore the HBV transcriptome. Furthermore, we developed Bolero, a computational method designed to handle the peculiarities of HBV sequencing data, which allows a detailed characterization of the HBV transcriptome.

## Introduction

Close to 290 million people worldwide are chronically infected with Hepatitis B virus (HBV) and are at high risk of developing severe liver diseases, such as cirrhosis and hepatocellular carcinoma [1]. Although prophylactic vaccines are available since 1981 and are effective in preventing HBV infection and transmission, current therapeutic options do not allow viral eradication in the large majority of patients. This stems mainly from the persistence of the HBV genome in the form of an episomal structure termed covalently closed circular (ccc)DNA [2]. Thus, developing novel therapies aimed at suppressing cccDNA transcriptional activity are of particular relevance. Moreover, the assessment of these molecules would also require the identification of reliable biomarkers that accurately reflect the intrahepatic HBV reservoir. In this context, recent reports support the usefulness of quantifying circulating HBV RNAs as an indicator of cccDNA activity [3]. However, the detailed characterization of such RNA species remains challenging due to the particular HBV biology. The HBV genome comprises a 3.2 kbp, circular, double-stranded DNA molecule with four distinct open reading frames, which are located on the negative strand and share a common transcription termination site (Fig 1). This genomic configuration serves as the sole template for eight primary RNAs, namely preCore, pgRNA, preS1, preS2, HBs, L-HBx, M-HBx, and S-HBx [4]. Interestingly, the longest RNAs, pgRNA and preCore, are 3.3 and 3.4 kb long respectively, exceeding the genome length. During their transcription that is initiated upstream of the termination site [4,5], the RNA polymerase II bypasses the termination site in the initial round and stops at the second passage, thereby creating 5’ and 3’ redundant extremities in these particular RNAs. Additionally, smaller RNAs exhibit substantial overlap, rendering them partially inclusive of each other regarding sequence content. Furthermore, twenty-two spliced variants have been identified so far, designated SP01 to SP22 [6]. Specifically, SP01 to SP20 arise from pgRNA splicing using both canonical and non-canonical splicing sites, while SP21 and SP22 stem from preS2/HBs RNAs. Given the considerable sequence similarity shared with the unspliced HBV RNAs and between each spliced variants, their discrimination relies on the identification of their respective transcription start site (TSS) and their splice junctions. Consequently, all HBV RNA sequences, including spliced variants, are subsumed under the longest HBV RNA sequence (i.e., preCore). A major advance in this field involved the development of 5’ rapid amplification of complementary DNA ends (5’RACE), as a strategy tailored to exclusively capture and amplify full-length HBV RNAs [5]. By employing 5’ end capping, mature transcripts are isolated and subsequently amplified via an HBV-specific primer located near the polyadenylation signal (PAS). This approach enables capturing canonical HBV transcripts alongside spliced variants derived from pgRNA and preS2/HBs RNAs. Associated to Sanger sequencing, this methodology allowed to discriminate all the major viral transcripts and demonstrate that they could be differentially regulated.

**Fig 1.**
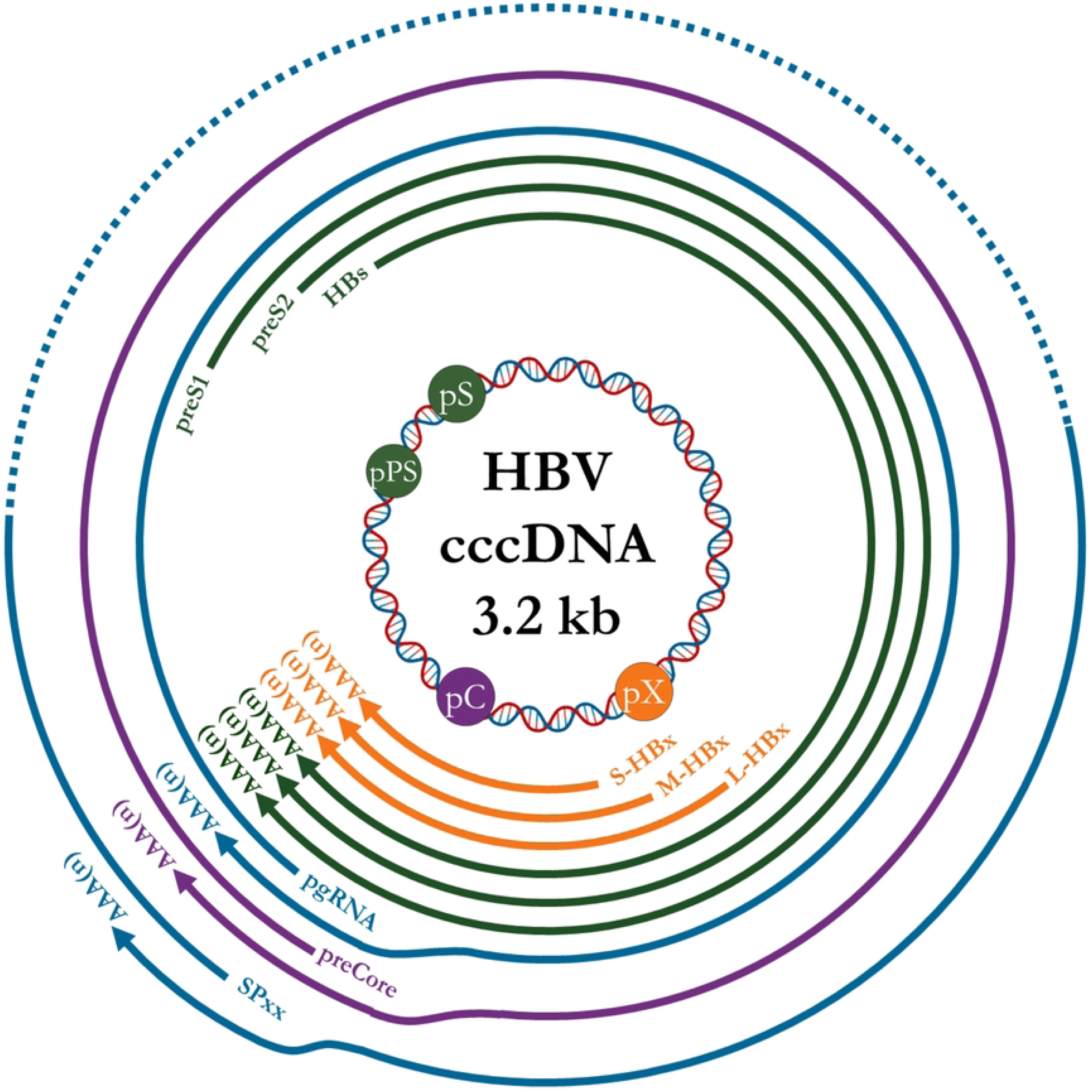
Schematic representation of HBV genome organization and viral transcripts. HBV cccDNA is represented in circular double stranded DNA form. Plain colored circles represent preCore/pg promoter (pC, purple), preS1 promoter (pPS, green), preS2/S promoter (pS, green) and HBx promoter (pX, orange). Circular lines represent the corresponding RNA species, preCore (purple), pgRNA (blue), preS1, preS2 and HBs (green), the three isoforms of HBx (L-HBx, M-HBx and S-HBx, orange). Dashed blue line represents spliced variants (SP01 to SP20, SP21 and SP22 are not represented). The “AAA(n)“ characters represent the polyA tail of HBV RNAs.

To investigate deeper, high throughput RNA-seq methodologies could be employed. However, due to the peculiar structure of the HBV genome, conventional short-read sequencing technologies failed in the identification and quantification of single HBV RNA molecules. Furthermore, the redundancy in the longest RNAs prevented the unique mapping of short reads originated from this particular region. Finally, a short-read uniquely mapped on a non-redundant region of the HBV RNAs sequences can be assigned to at least two different RNAs because of their massive overlapping. More recently, long-read and full-length molecule sequencing technologies, such as Oxford Nanopore Technology (ONT), allow to sequence full-length nucleotidic molecules [7]. Therefore, we took advantage of this approach and coupled the 5’RACE method [5] to capture and amplify HBV transcripts to the ONT long-read sequencing method, as a means to obtain full-length HBV transcript sequences. To be analyzed, such sequencing data require nonetheless a dedicated bioinformatics pipeline that is not available yet. We thus developed Bolero, a bioinformatics method and built-in workflow to evaluate the relative expression of HBV RNAs and their spliced variants, named after the theme composed by Maurice Ravel in which the successive entry of instruments creates an overlapping structure resembling that of HBV transcripts. The filtering and analyses steps in the Bolero pipeline allow to assign 99% of the preprocessed reads to each particular HBV RNA species, enabling their relative levels to be represented as percentage of global viral genes expression and thus, giving the composition of the HBV transcriptome in a given sample (Fig 2). Moreover, this pipeline stores information of putative TSS associated with particular splice junctions for each read, allowing the potential identification of novel spliced variants. Therefore, Bolero represents a useful tool for the detailed characterization of the HBV transcriptome.

**Fig 2.**
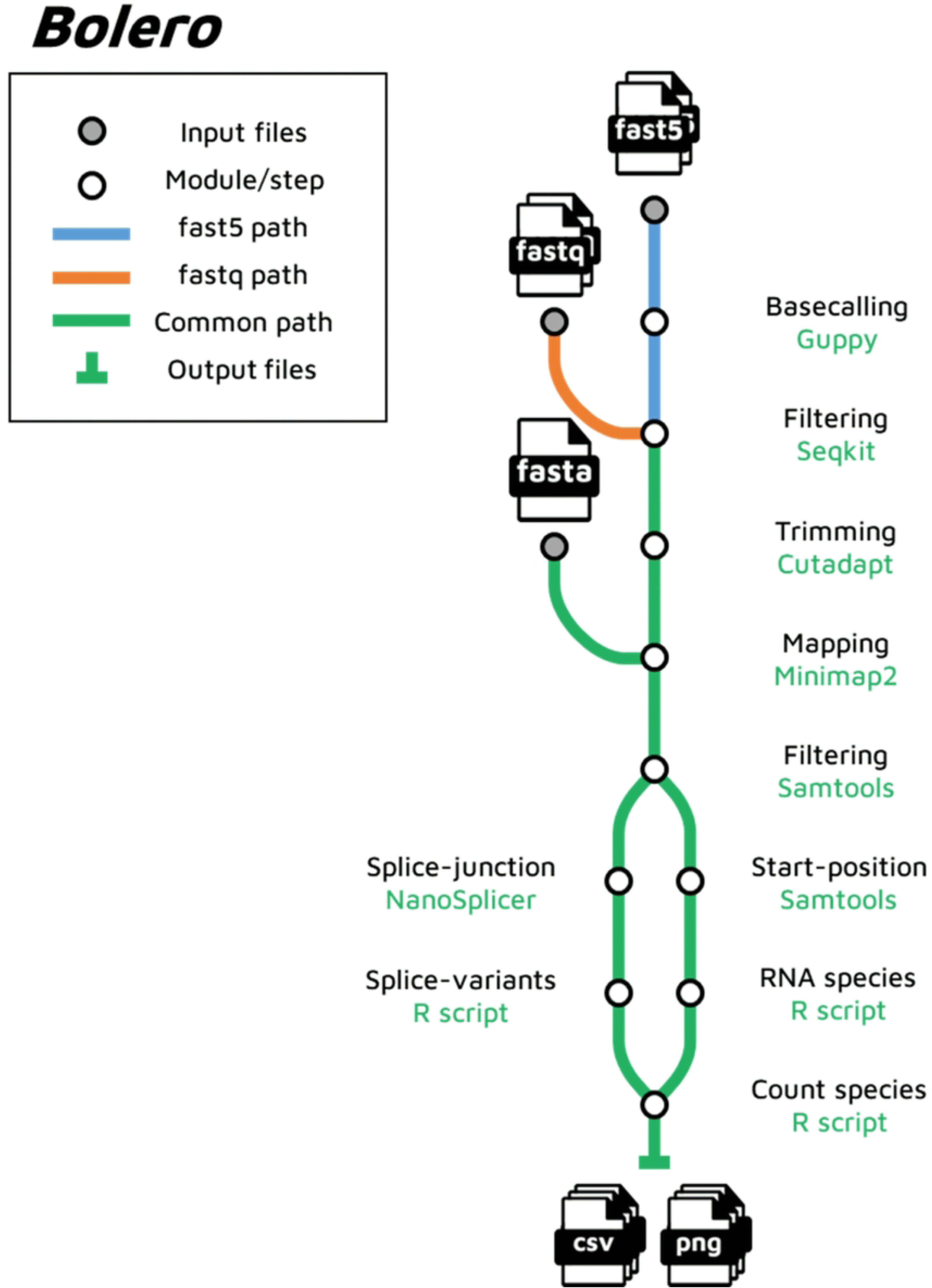
Overview of the Bolero workflow. Metro map visualization of the Bolero workflow for processing Nanopore long-read sequencing data from 5’RACE-amplified HBV RNAs. Input files include either raw Fast5 files (blue line) or basecalled Fastq files (orange line); subsequent stages follow the green line regardless of input format. Stages requiring additional inputs are depicted as grey stations, with white stations indicating the various steps of the workflow, denoted by the step name in black and the corresponding program in green. The final station marks the endpoint of the workflow, producing output files.

## Results

### Filtering of raw data: removing reads that do not correspond to full-length HBV RNAs

To develop and adapt the different steps of the workflow, triplicate synthetic datasets were simulated as toy datasets. Each dataset contained 2.5×10^5^ reads per HBV RNA species. The reads were designed to be similar to the expected ones produced by Nanopore sequencing of 5’RACE-amplified products, consisting of quasi full-length HBV RNAs linked to the 5’RACE adapter sequence. In order to consider only nearly full-length HBV sequenced RNAs captured by the 5’RACE-PCR method, reads underwent several filtering steps. Initially, reads that did not contain both the 5’RACE adapter and the gene specific primer (GSP) sequences were excluded from the analysis. To identify reads corresponding to HBV RNAs using the TSS position on the reference genome, only mature and capped RNAs should be considered. Hence, reads were not trimmed, except for the 5’RACE and Nanopore technical adapters that did not correspond to HBV RNAs sequence. The number of post-filtering reads was found to depend on both the sequencing yield and the efficiency of the 5’RACE procedure in capturing complete HBV RNA transcripts. In order to evaluate and optimize each stage of the workflow, synthetic datasets composed of simulated reads were employed. Specifically, these synthetic datasets contained on average 11% of synthetic reads with the 5’RACE adapter sequence, 4% harbouring the GSP sequence, and 4% exhibiting lengths greater than 390 bp. Upon applying filtering measures to this synthetic dataset, only 4% of the initial synthetic reads remained eligible for further investigation. This corresponded to a total mean quantity of 323,535 filtered reads derived from an input pool of 7.5×10^6^ raw synthetic reads per dataset (Table 1).

**Table 1.**
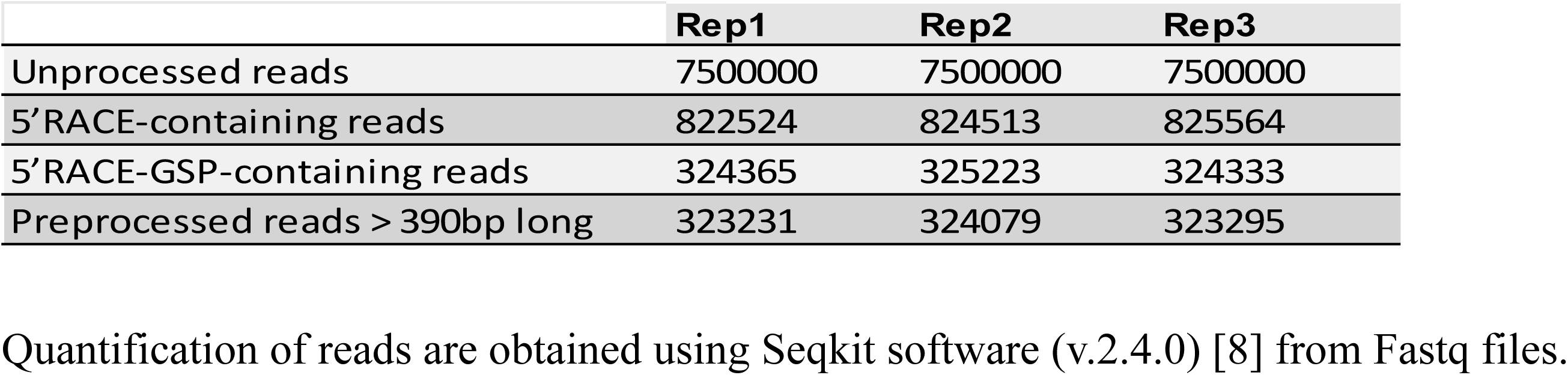
Quantification of synthetic reads following successive filtering steps.

### HBV reference sequence and mapping: HBV full-length RNAs require a longer reference sequence than the linearized HBV genome itself

Given the fact that the HBV genome is circular and the longest transcripts are longer than the genome itself, the choice of reference sequence is crucial. As the preCore RNA encompasses all the known HBV transcripts, including the spliced variants, we decided to use this sequence as reference to map the filtered reads. This sequence is defined from the preCore TSS (position 1692 from the EcoRI site) to the PAS (position 1932) along 1.1X genome of the HBV genotype D subtype ayw, which is 3,422 bp in length [9]. Minimap2, a cutting-edge software tool for aligning long-read sequencing data, was selected for the mapping step of the workflow due to 1) its preset mode for ONT reads, 2) its awareness of splicing events, and 3) its ability to detect non-canonical splicing sites [10,11]. These features are particularly relevant given the presence of previously described HBV spliced variants exhibiting non-canonical splicing sites [6]. After preprocessing and filtering, reads were mapped to the reference HBV sequence. Considering that each filtered read corresponded to a single RNA molecule proportionally amplified by the 5’RACE method, only the best alignment for each read was retained, and secondary alignments were disregarded to prevent potential overestimation of expression levels. Out of the filtered and preprocessed reads previously acquired, almost all exhibited unique mappings to the preCore reference sequence and all possessed a FLAG value of “0” (Table 2). The very small number of unmapped reads corresponded to highly noisy synthetic reads.

**Table 2.** Mapped and unmapped synthetic HBV read quantities.

|  | Rep1 | Rep2 | Rep3 |
| --- | --- | --- | --- |
| Input reads | 323 231 | 324 079 | 323 295 |
| **Mapped reads** | **323196** | **324046** | **323246** |
| Unmapped reads | 35 | 33 | 49 |
| **% mapped reads** | **99.99%** | **99.99%** | **99.98%** |

Quantification of mapped and unmapped reads are obtained using samtools (v.1.17) [12,13].

### Start position extraction: Sequence specificity is not sufficient for the identification of HBV RNAs

Each HBV RNA species differs from each other by their unique TSS position. Thus, as a first step, we extracted the TSS position of each HBV RNA species we had previously identified by Sanger sequencing [5]. Taking the EcoRI restriction site as the position 1, we were able to identify the TSS coordinates for the preCore (1692), pgRNA (1818), preS1 (2805), preS2 (3157), HBs (7), L-(1065), M-(1243) and S-HBx (1418) RNAs (Table 3). To assign the reads to their corresponding HBV RNA species, we then identified the start position of the alignment for each read. This position, corresponding to the first nucleotide aligned to the reference sequence, is directly extracted from the BAM file and corresponds to the POS field [12]. Given that errors are known to be made by ONT sequencing, especially at 5’ ends [14], we expect the reads start position mapped to the reference sequence to vary around the theoretical TSS of the corresponding RNA species. To define intervals of the starting position of every aligned read, we used simulated triplicated datasets containing the sequence of individual HBV RNAs with their respective TSS and determined the range of start position for each simulated read. As shown in Fig 3, for the three simulated datasets containing only pgRNA reads, start position distributions are very similar and comprised between 1799 (in the rep3) and 1904 (in the rep2). To capture as much information as possible from these reads, we considered the largest range taking into account the minimum and maximum start positions per HBV RNA species through the three simulated datasets (Table 3). Therefore, these defined start position ranges determine putative TSS assigned to each read based on the start position of its alignment.

**Fig 3.**
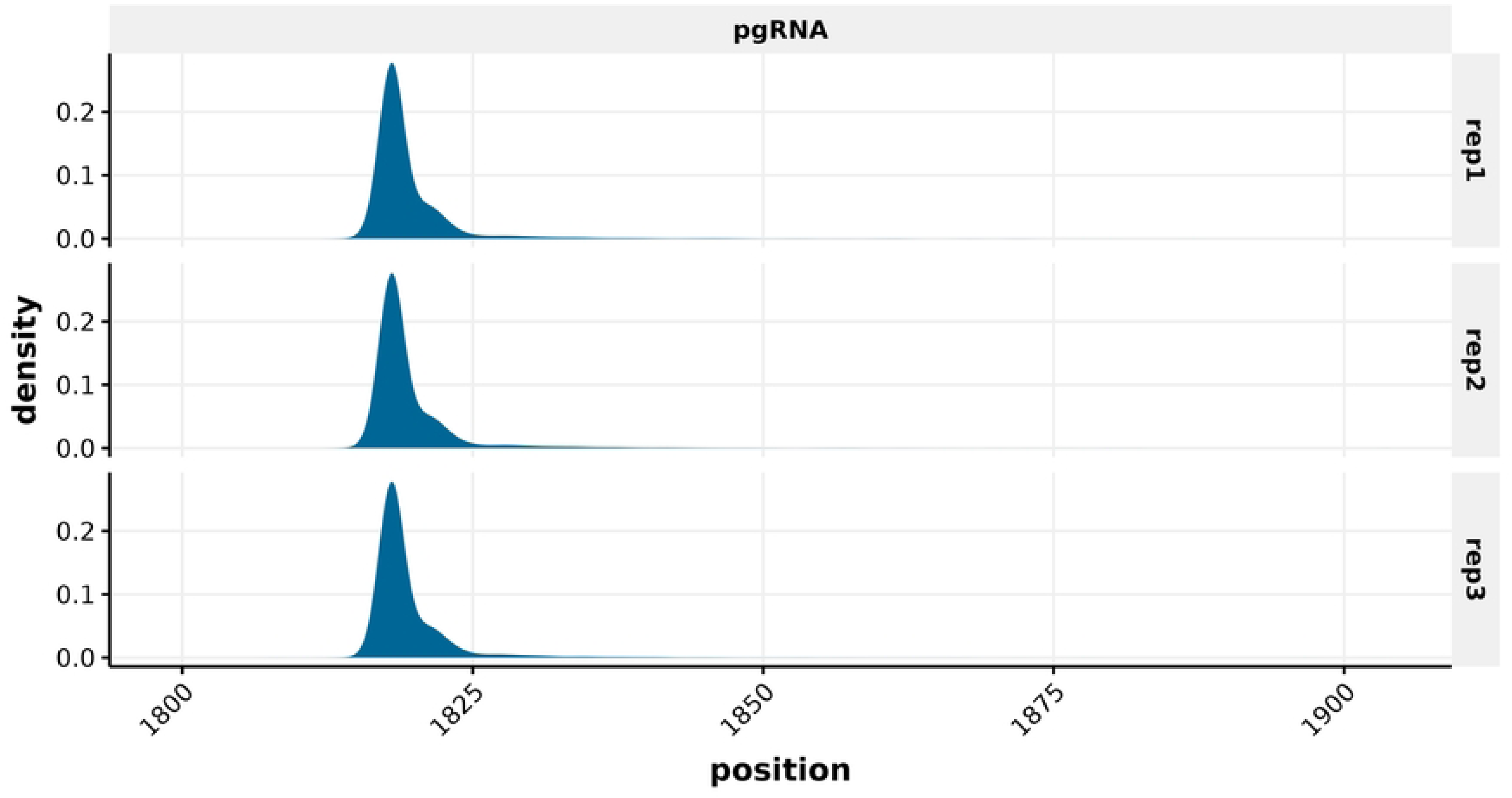
Start position distribution of pgRNA-derived reads in triplicate synthetic datasets. Positions are EcoRI based on the HBV genome genotype D, strain ayw.

**Table 3.** Minimum and maximum start positions, theoretical TSS positions and selected borders employed to define putative TSS of HBV reads.

| Species | Theoretical TSS position | Rep1 |  | Rep2 |  | Rep3 |  | Selected borders |  |  |
| --- | --- | --- | --- | --- | --- | --- | --- | --- | --- | --- |
|  |  | Min | Max | Min | Max | Min | Max | Min | Max | Range |
| preCore | 1692 | 1692 | 1798 | 1692 | 1750 | 1692 | 1763 | **1692** | **1798** | **1692-1798** |
| pgRNA | 1818 | 1814 | 1887 | 1814 | 1904 | 1799 | 1903 | **1799** | **1904** | **1799-1904** |
| preS1 | 2805 | 2798 | 2937 | 2798 | 2909 | 2798 | 2940 | **2798** | **2940** | **2798-2940** |
| preS2 | 3157 | 3146 | 51 | 3147 | 67 | 3146 | 63 | **3146** | **67** | **3146-175** |
| HBs | 7 | 3168 | 175 | 3171 | 72 | 3168 | 68 | **3168** | **175** |  |
| L_HBx | 1065 | 1062 | 1154 | 1062 | 1137 | 1062 | 1140 | **1062** | **1154** | **1062-1154** |
| M_HBx | 1243 | 1236 | 1349 | 1237 | 1326 | 1236 | 1366 | **1236** | **1366** | **1236-1366** |
| S_HBx | 1418 | 1407 | 1493 | 1409 | 1519 | 1411 | 1510 | **1407** | **1519** | **1407-1519** |
| HBx | - | - | - | - | - | - | - | **1062** | **1519** | **1062-1519** |

In synthetic datasets, most reads align with their starting positions matching the theoretical TSS of the corresponding RNA species (Fig 4 and S1-S3 Figs). The start position intervals for pgRNA (Fig 4A, B and S4 Fig), preCore (Fig 4B and S4 Fig) and preS1 (S1-S3 Figs), as well as different isoforms of HBx (Fig 4C and S5 Fig) differ from those of other RNAs, allowing for unequivocal assignment. However, the situation is more complex for preS2 and HBs (Fig 4D and S6 Fig). Indeed, preS2 and HBs have overlapping start position intervals, which represents an obstacle to unequivocally assign reads to specific RNA species. Thus, in order to minimize this ambiguity when classifying reads, we grouped together all reads whose start positions fall within the range of 3146 to 175 under the category of preS2/S RNAs, without distinguishing between them (Table 3). Then, we combined all datasets into a single one containing simulated reads corresponding to every individual HBV RNA species to estimate their relative quantity based on the start position ranges we had previously determined (Fig 4E and S7 Fig). Relative quantification of reads per start position ranges corresponds to the groups previously defined (Fig 5A and S8 Fig). As anticipated, since we established start position ranges for each HBV RNA species based on the same dataset, all simulated reads were allocated to their appropriate categories (Fig 5B, Table 4 and S8 Fig). In line with this, the corresponding spliced variants were also accurately classified. Specifically, spliced variants SP01 to SP20 were attributed to the pgRNA group, whereas spliced variants SP21 and SP22 fell under the preS2/S group. As a result, a putative TSS is assigned to each read.

**Fig 4.**
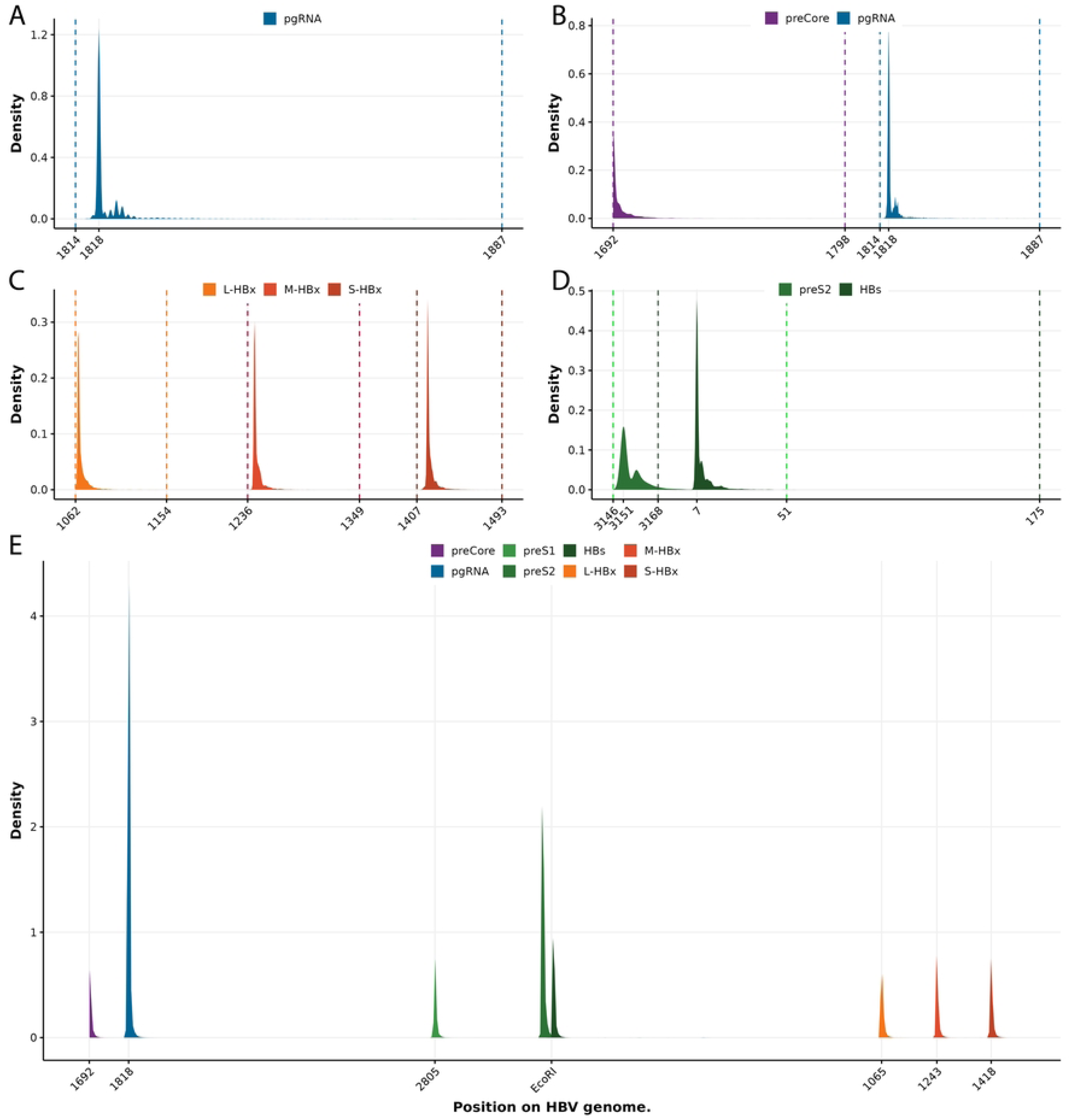
Start positions distribution in synthetic datasets (replicate 1). Positions are EcoRI based on the HBV genome genotype D strain ayw. (A-D) Start position distributions for distinct HBV RNA species: (A) pgRNA, (B) preCore + pgRNA, (C) three isoforms of HBx (L-, M-, and S-HBx), and (D) preS2 + HBs. Vertical dashed lines denote minimum and maximum start positions for each respective HBV RNA species. (E) Overall start position distribution for merged synthetic dataset reads. All distributions are presented as percentages of total reads for each panel separately.

**Fig 5.**
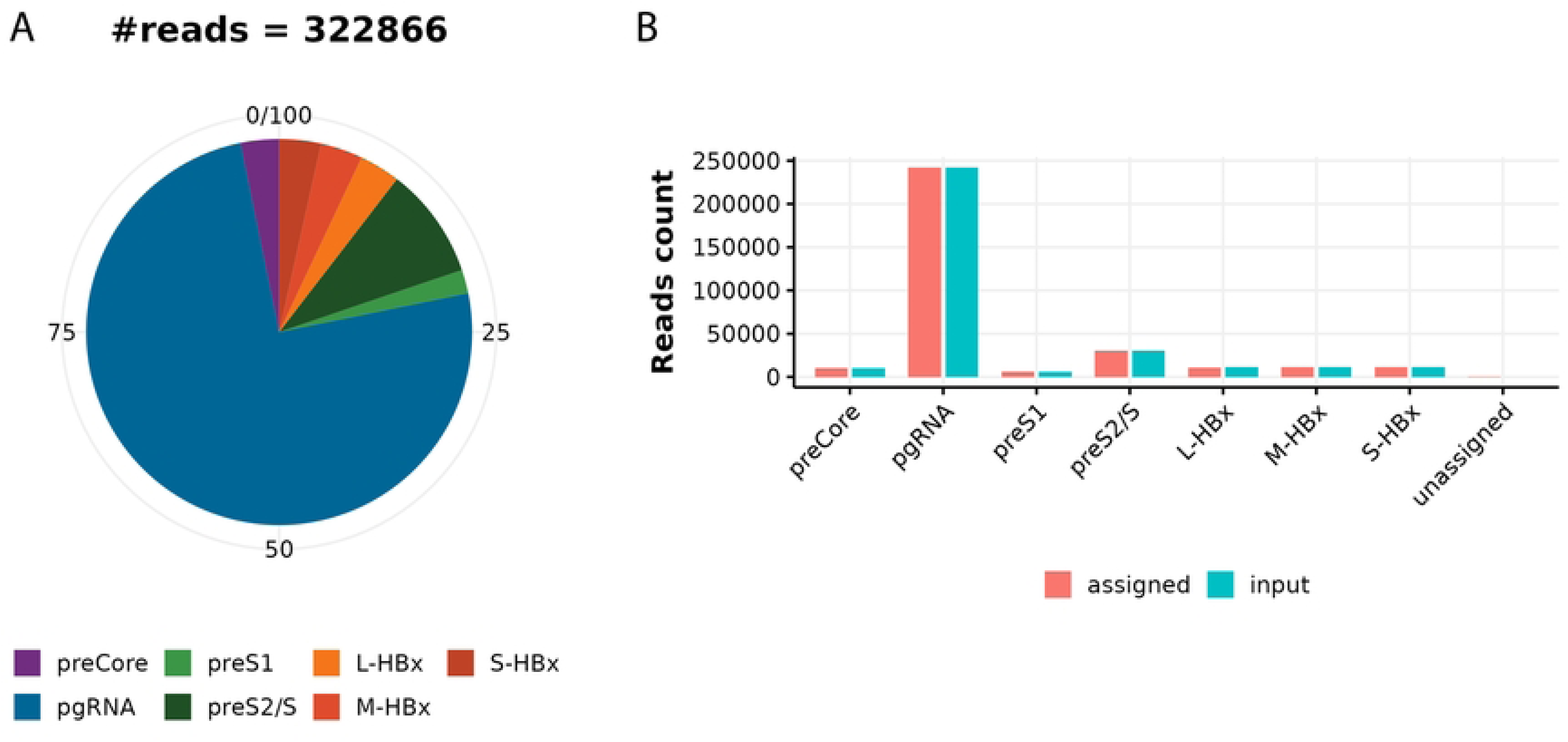
Relative and absolute read quantification according to putative TSS groups in synthetic datasets (replicate 1). (A) Pie chart depicting relative quantities of reads assigned to putative TSS groups. #reads represents the total number of assigned reads. (B) Comparison between absolute quantities of reads assigned to a putative TSS by Bolero based on start positions and preprocessed simulated reads as input of quantification step.

**Table 4.** Quantity of input reads compared to those assigned with Bolero for each individual HBV RNAs dataset based on putative TSS recognition.

| TSS | Rep1 |  | Rep2 |  | Rep3 |  |
| --- | --- | --- | --- | --- | --- | --- |
|  | input | assigned | input | assigned | input | assigned |
| L-HBx | 11229 | 11066 | 11325 | 11182 | 11013 | 10865 |
| M-HBx | 11399 | 11399 | 11348 | 11348 | 11303 | 11303 |
| S-HBx | 11284 | 11284 | 11455 | 11454 | 11346 | 11346 |
| pgRNA | 242337 | 242175 | 242408 | 242273 | 242330 | 242199 |
| preCore | 10386 | 10385 | 10460 | 10460 | 10327 | 10327 |
| preS1 | 6395 | 6398 | 6375 | 6376 | 6367 | 6371 |
| preS2/S | 30166 | 30159 | 30675 | 30671 | 30560 | 30555 |
| total | 323196 | 322866 | 324046 | 323764 | 323246 | 322966 |

Quantification of input and assigned reads are obtained using Bolero R scripts in three independent experiments.

### Spliced variant identification and quantification

The majority of reads are grouped into specific RNA species categories based on their start positions or, more broadly, their putative TSS positions. Some of these reads may include splice junctions. Distinguishing known spliced variants, as documented in the literature, is possible by considering two distinct characteristics: their 1) TSS position and 2) unique combination of splice junctions. To identify spliced variants from the reads, we employed the Nanosplicer software [15]. Nanosplicer integrates a feature that identifies splice junctions in each read, with those lacking such junctions falling into the canonical RNA species categories. Conversely, reads containing splice junctions are scrutinized and assigned to the corresponding spliced variants (i.e., SP01 to SP22), taking into consideration both their start positions and the specific combinations of splice junctions they contain. However, similar to the considerations for start positions, it is necessary to define position intervals for splice junction sites. This entails analysing simulated and independent spliced variant datasets to determine the position ranges of donor and acceptor sites. This is based on triplicate simulated datasets of spliced variants containing unique splice junctions. Some spliced variants share the same splice junctions; thus, they were named based on their positions (Fig 6 and Table 5). For example, J2450_490 is the unique junction of SP01 and is shared by SP02, SP04, and SP15, whereas J2070_490 is uniquely present in the SP03 spliced variant.

**Fig 6.**
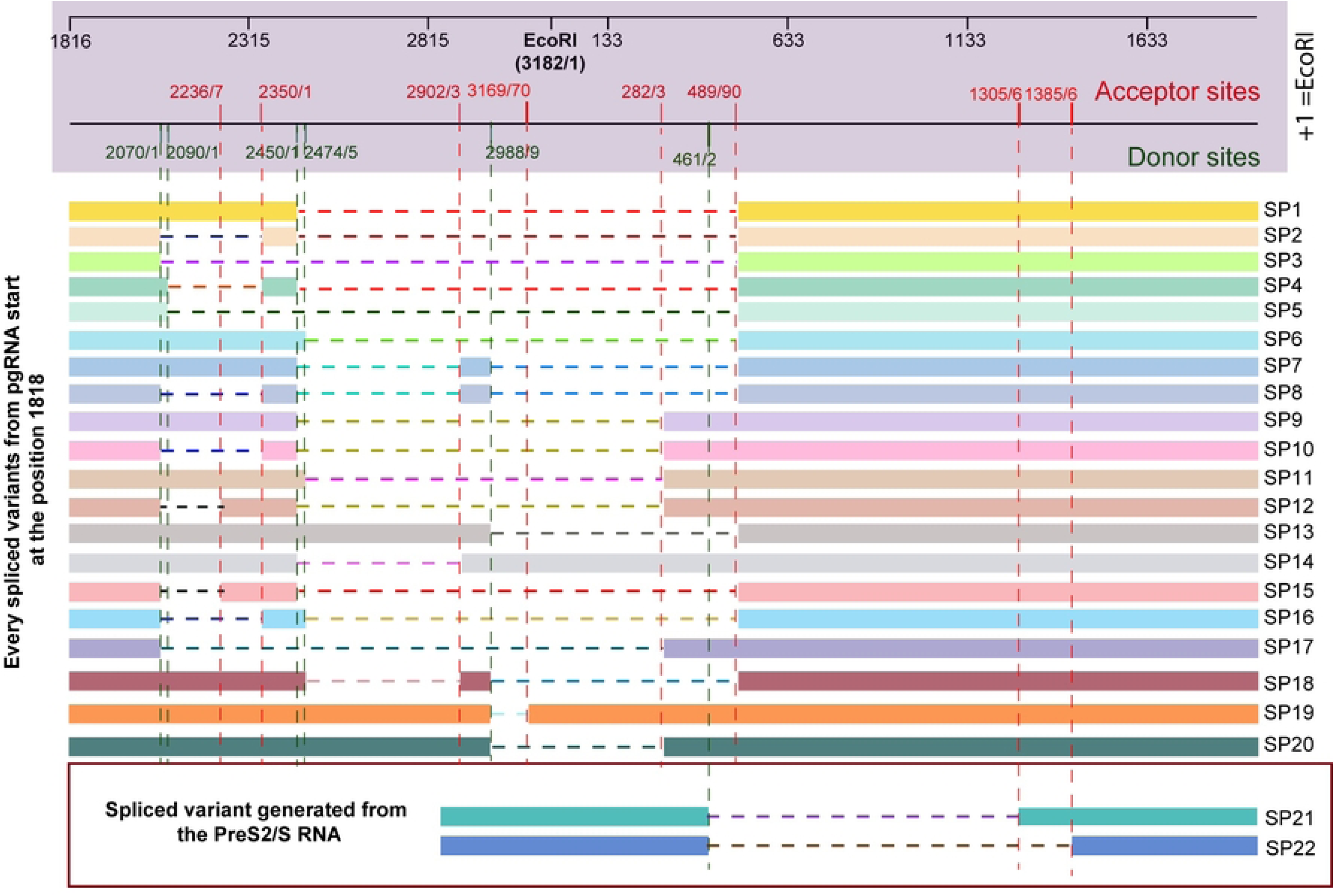
Schematic representation of HBV spliced variants (SP01 to SP22). Positions are EcoRI based on the HBV genome genotype D, strain ayw.

**Table 5.**
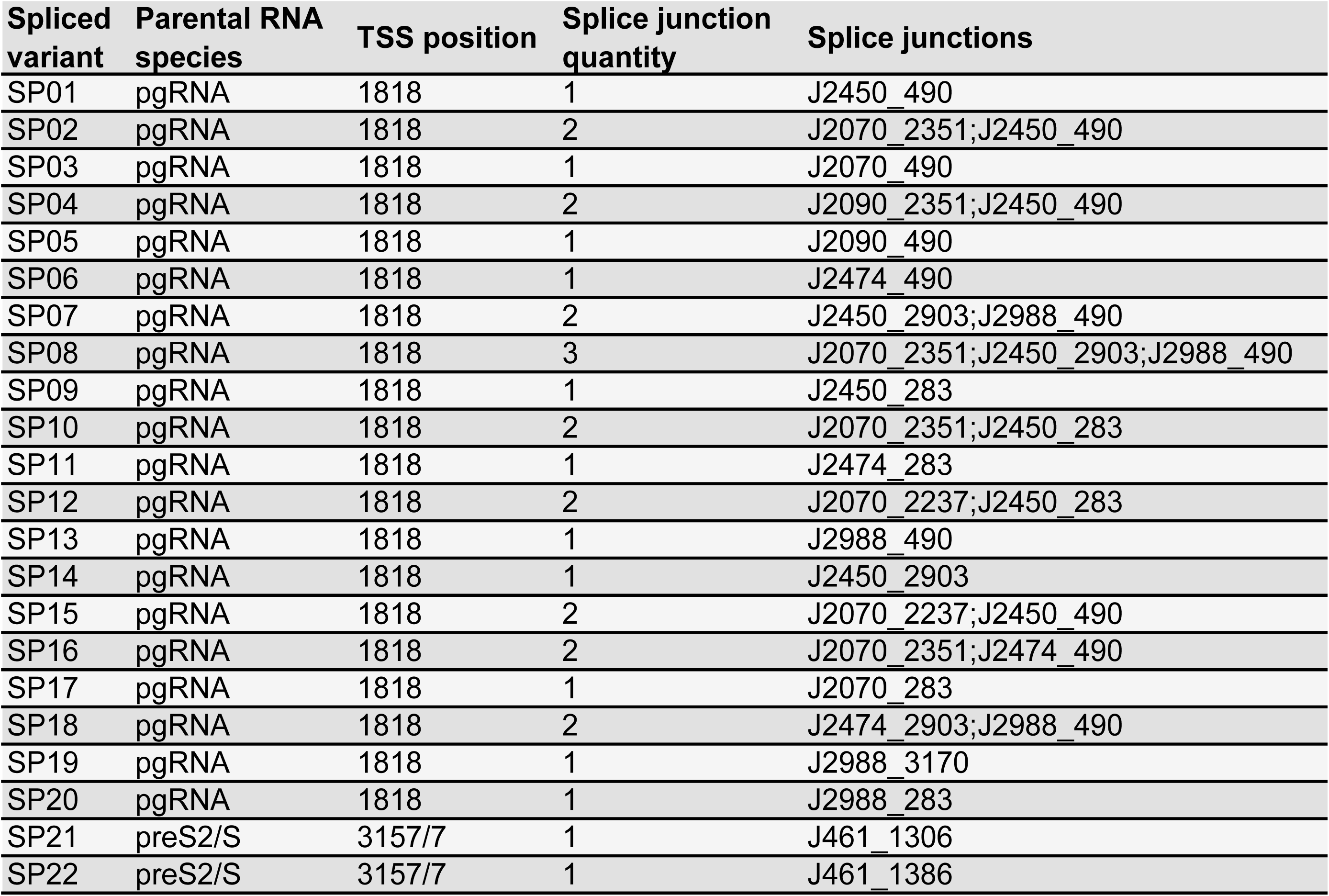
Description of HBV spliced variants.

Analyzing simulated datasets of spliced variants, we assessed the distribution of donor and acceptor site positions as determined by Nanosplicer (Fig 7 and S9 Fig). Upon closer examination of the donor and acceptor site position distributions of the J2450_490 junction (Fig 7A, B), we were able to determine the position ranges. We observed that the minimum and maximum positions of the J2450_490 donor site (position range: 2424-2482) and the J2474_283 donor site (position range: 2437-2519) overlapped (Fig 7C and S9 Fig). After careful consideration, we excluded extreme values (i.e., quantiles 0.005 and 0.995) accounting for positions representing 99.9% of reads from respective spliced variants. As a result, we established distinct position ranges for each donor and acceptor site (Table 6).

**Fig 7.**
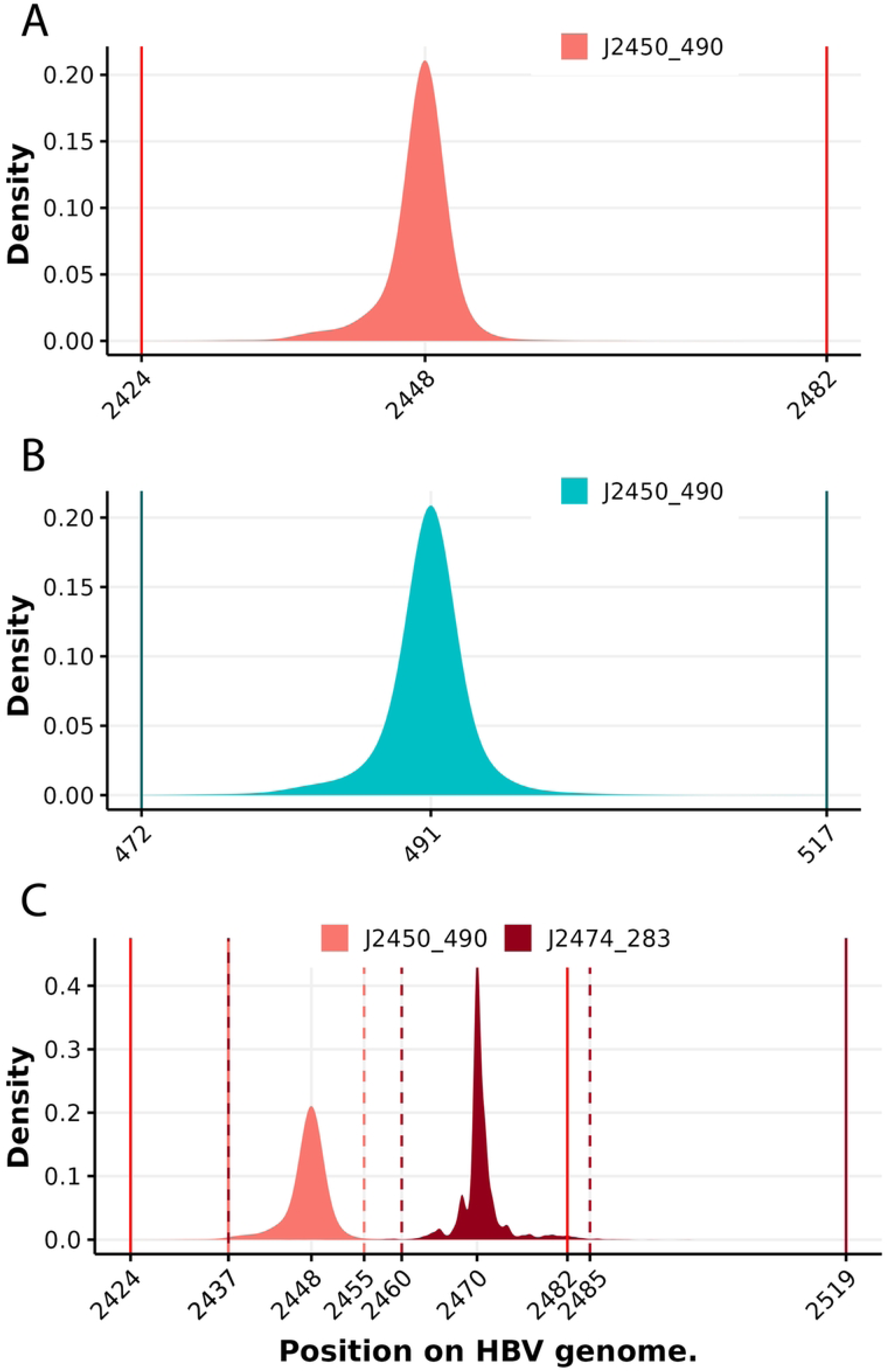
Density of reads expressed as a function of donor (red) or acceptor (blue) sites positions of splice junctions. (A, B) Position distributions of the (A) donor and (B) acceptor sites from junction J2450_490. (C) Position distributions of donor sites from J2450_490 and J2474_283 junctions. Plain vertical lines represent minimum and maximum positions, dashed vertical lines indicate selected borders, defining the interval containing 99.9% of donor or acceptor site positions.

**Table 6.** Description of HBV splice junctions.

| Splice junction | Donor site position | Acceptor site position | Donor site position range | Acceptor site position range | Spliced variants | Junction sequence |
| --- | --- | --- | --- | --- | --- | --- |
| J461_1306 | 461 | 1306 | 447-469 | 1297-1314 | SP21 | ACTATCAAG//CAGGTCTGG |
| J461_1386 | 461 | 1386 | 447-469 | 1377-1396 | SP22 | ACTATCAAG//CTGTGCTGC |
| J2070_2237 | 2070 | 2237 | 2056-2075 | 2229-2246 | SP12;SP15 | TGCACTCAG//AGAAACCGT |
| J2070_2351 | 2070 | 2351 | 2056-2075 | 2341-2360 | SP02;SP08;SP10;SP16 | TGCACTCAG//ACGACGAGG |
| J2070_283 | 2070 | 283 | 2056-2075 | 274-297 | SP17 | TGCACTCAG//GGGGAAC TA |
| J2070_490 | 2070 | 490 | 2056-2075 | 478-499 | SP03 | TGCACTCAG//GATCCTCAA |
| J2090_2351 | 2090 | 2351 | 2077-2096 | 2341-2360 | SP04 | TTTGCTGGG//ACGACGAGG |
| J2090_490 | 2090 | 490 | 2077-2096 | 478-499 | SP05 | TTTGCTGGG//GATCCTCAA |
| J2450_283 | 2450 | 283 | 2436-2457 | 274-297 | SP09;SP10;SP12 | AACCTCAAT//GGGGAAC TA |
| J2450_2903 | 2450 | 2903 | 2436-2457 | 2893-2912 | SP07;SP08;SP14 | AACCTCAAT//TTGGATCCA |
| J2450_490 | 2450 | 490 | 2436-2457 | 478-499 | SP01;SP02;SP04;SP15 | AACCTCAAT//GATCCTCAA |
| J2474_283 | 2474 | 283 | 2460-2487 | 274-297 | SP11 | ACTCATAAG//GGGGAAC TA |
| J2474_2903 | 2474 | 2903 | 2460-2487 | 2893-2912 | SP18 | ACTCATAAG//TTGGATCCA |
| J2474_490 | 2474 | 490 | 2460-2487 | 478-499 | SP06;SP16 | ACTCATAAG//GATCCTCAA |
| J2988_283 | 2988 | 283 | 2976-2995 | 274-297 | SP20 | GCCAACAAG//GGGGAAC TA |
| J2988_3170 | 2988 | 3170 | 2976-2995 | 3161-3179 | SP19 | GCCAACAAG//CCATGCAGT |
| J2988_490 | 2988 | 490 | 2976-2995 | 478-499 | SP07;SP08;SP13;SP18 | GCCAACAAG//GATCCTCAA |

Finally, the combination of start positions and splice junctions allows to unequivocally identify each spliced variant and discriminate them from their unspliced parental RNA. As an example, a read whose start position is in the range of pgRNA TSS and containing the J2070_490 junction will be assigned to the SP03 variant, while another read with the same start position without splice junction will be assigned to pgRNA. In some particular cases, start positions and splice junctions do not correspond to previously reported ones, thus, the read cannot be assigned to an HBV RNA species and will be tagged as “Undefined”. This is the case for 3090 reads in the complete simulated dataset rep1 (Fig 8, Table 7 and S10 Fig), representing 0.96% of error. This error is mainly due to the 0.1% of reads that are excluded to precisely define splice junction position ranges. As expected, the quantity of simulated reads is comparable to the quantification obtained using Bolero (Fig 8, Table 7 and S10 Fig). By combining start position and splice junction identification, we are able to assign 99% of the mapped reads to the different known HBV RNA species. As a result, relative levels of HBV RNAs are obtained as a percentage of global viral genes expression, thus giving a composition of the HBV transcriptome in the sample (Fig 9A, B and S11 Fig).

**Fig 8.**
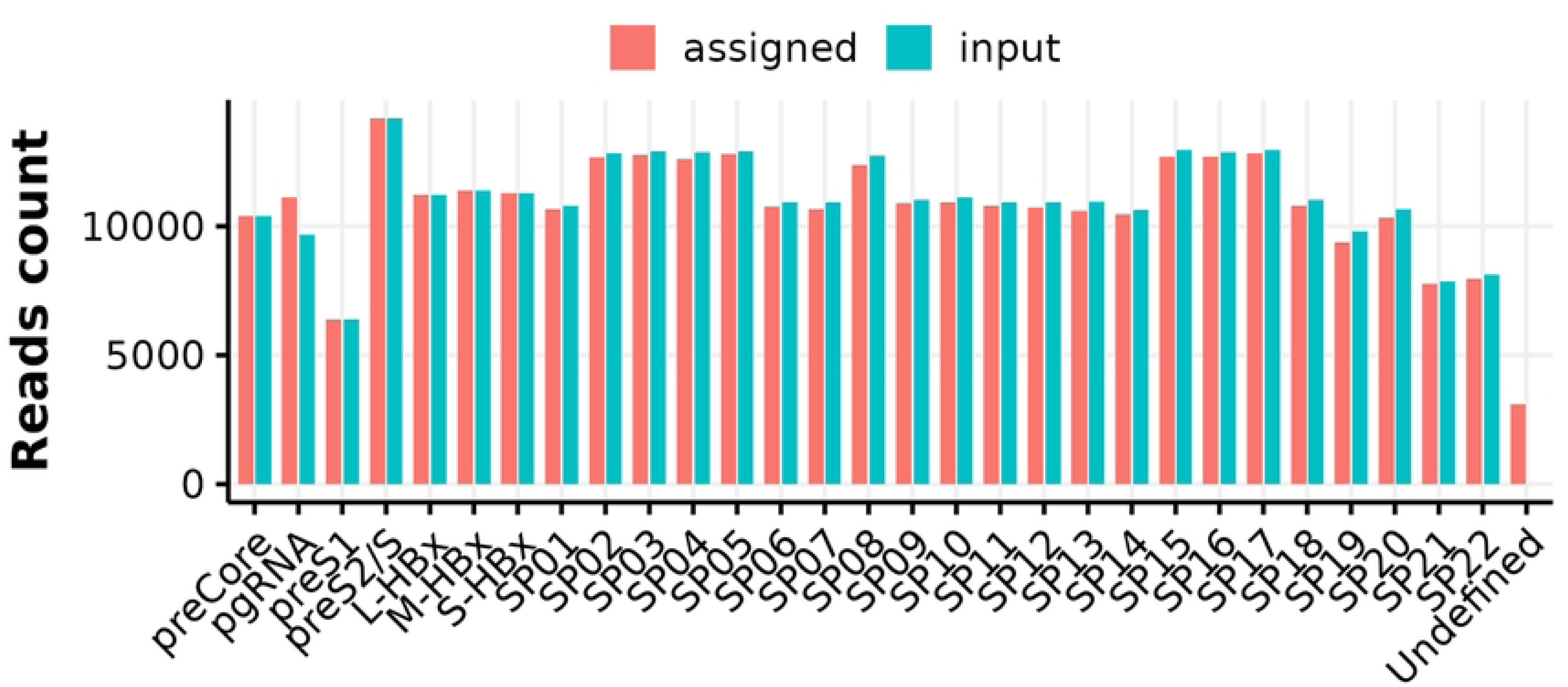
Comparison between absolute quantities of reads assigned by Bolero and preprocessed simulated reads as input of the quantification step in synthetic datasets (replicate 1). Quantification takes into account the start position of reads and the presence of splice junctions.

**Fig 9.**
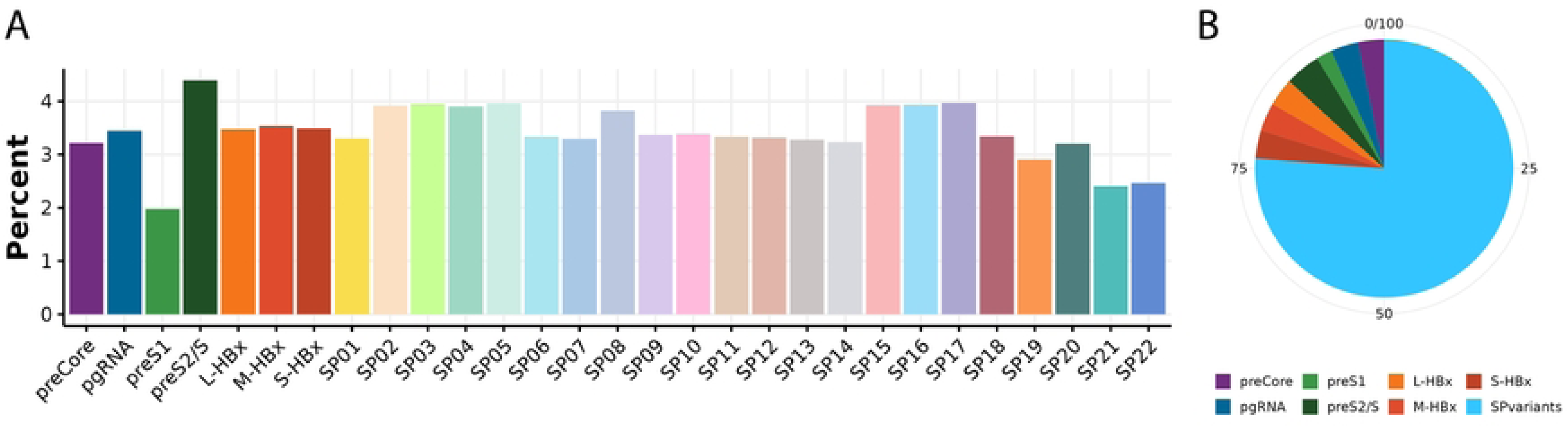
Relative expression of HBV RNA species in synthetic datasets (replicate 1). (A) Bar chart of relative quantification of known HBV RNA species. (B) Pie chart of relative quantification of known canonical RNA species and grouped spliced variants.

**Table 7.** Quantity of input reads compared to those assigned with Bolero for each individual HBV RNAs dataset based on putative TSS and splice junctions’ composition.

| Species | Rep1 |  | Rep2 |  | Rep3 |  |
| --- | --- | --- | --- | --- | --- | --- |
|  | Input | Assigned | Input | Assigned | Input | Assigned |
| preCore | 10386 | 10385 | 10460 | 10460 | 10327 | 10327 |
| pgRNA | 9672 | 11129 | 9644 | 11047 | 9578 | 11017 |
| preS1 | 6395 | 6394 | 6375 | 6375 | 6367 | 6366 |
| preS2/S | 14189 | 14188 | 14445 | 14445 | 14346 | 14346 |
| L-HBx | 11229 | 11229 | 11325 | 11325 | 11013 | 11013 |
| M-HBx | 11399 | 11399 | 11348 | 11348 | 11303 | 11303 |
| S-HBx | 11284 | 11282 | 11455 | 11450 | 11346 | 11344 |
| SP01 | 10780 | 10664 | 10977 | 10855 | 10890 | 10752 |
| SP02 | 12826 | 12656 | 12702 | 12565 | 12809 | 12636 |
| SP03 | 12888 | 12767 | 12894 | 12772 | 12933 | 12835 |
| SP04 | 12879 | 12612 | 13144 | 12875 | 12855 | 12569 |
| SP05 | 12894 | 12806 | 12871 | 12771 | 12892 | 12790 |
| SP06 | 10911 | 10773 | 11028 | 10902 | 11100 | 10971 |
| SP07 | 10929 | 10643 | 11118 | 10813 | 10839 | 10577 |
| SP08 | 12725 | 12355 | 12805 | 12409 | 12925 | 12514 |
| SP09 | 11023 | 10887 | 10905 | 10762 | 10913 | 10803 |
| SP10 | 11113 | 10935 | 11113 | 10939 | 11185 | 11007 |
| SP11 | 10916 | 10775 | 10920 | 10780 | 10733 | 10571 |
| SP12 | 10909 | 10719 | 10972 | 10775 | 11018 | 10827 |
| SP13 | 10964 | 10592 | 10867 | 10535 | 10774 | 10417 |
| SP14 | 10626 | 10453 | 10708 | 10538 | 10778 | 10605 |
| SP15 | 12959 | 12685 | 12736 | 12496 | 12751 | 12511 |
| SP16 | 12870 | 12704 | 12705 | 12525 | 13029 | 12846 |
| SP17 | 12976 | 12843 | 12938 | 12795 | 12880 | 12764 |
| SP18 | 11015 | 10787 | 10896 | 10661 | 11064 | 10818 |
| SP19 | 9810 | 9368 | 9767 | 9348 | 9750 | 9336 |
| SP20 | 10652 | 10342 | 10698 | 10393 | 10634 | 10311 |
| SP21 | 7858 | 7772 | 8115 | 8045 | 8122 | 8036 |
| SP22 | 8119 | 7962 | 8115 | 7939 | 8092 | 7911 |
| Total | 323196 | 320106 | 324046 | 320943 | 323246 | 320123 |
| Undefined | 0 | 3090 | 0 | 3103 | 0 | 3123 |

Quantification of input and assigned reads are obtained using Bolero R scripts in three independent replicates.

### Bolero validation: Quantification of RNA species in HBV-infected HepG2-NTCP cells

The above-described analyses clearly demonstrate that Bolero is efficient in identifying and quantifying HBV RNA species from simulated datasets. Thus, as a second step, we determined whether Bolero was also efficient in properly evaluating the HBV transcriptome in datasets generated from biological samples. With this aim, we performed HBV-specific 5’RACE-PCR experiments on RNAs extracted from three independent replicates of HBV-infected HepG2-NTCP cells. Amplicons were then sequenced using ONT and the raw sequencing data were processed and analysed using Bolero. The global pattern of start positions, expressed as percentage of total reads per sample, revealed a strong reproducibility between the three replicates (Fig 10A). Indeed, five main peaks, corresponding to the major HBV RNA species start positions, were identified close to the regions corresponding to HBV RNAs TSS locations.

**Fig 10.**
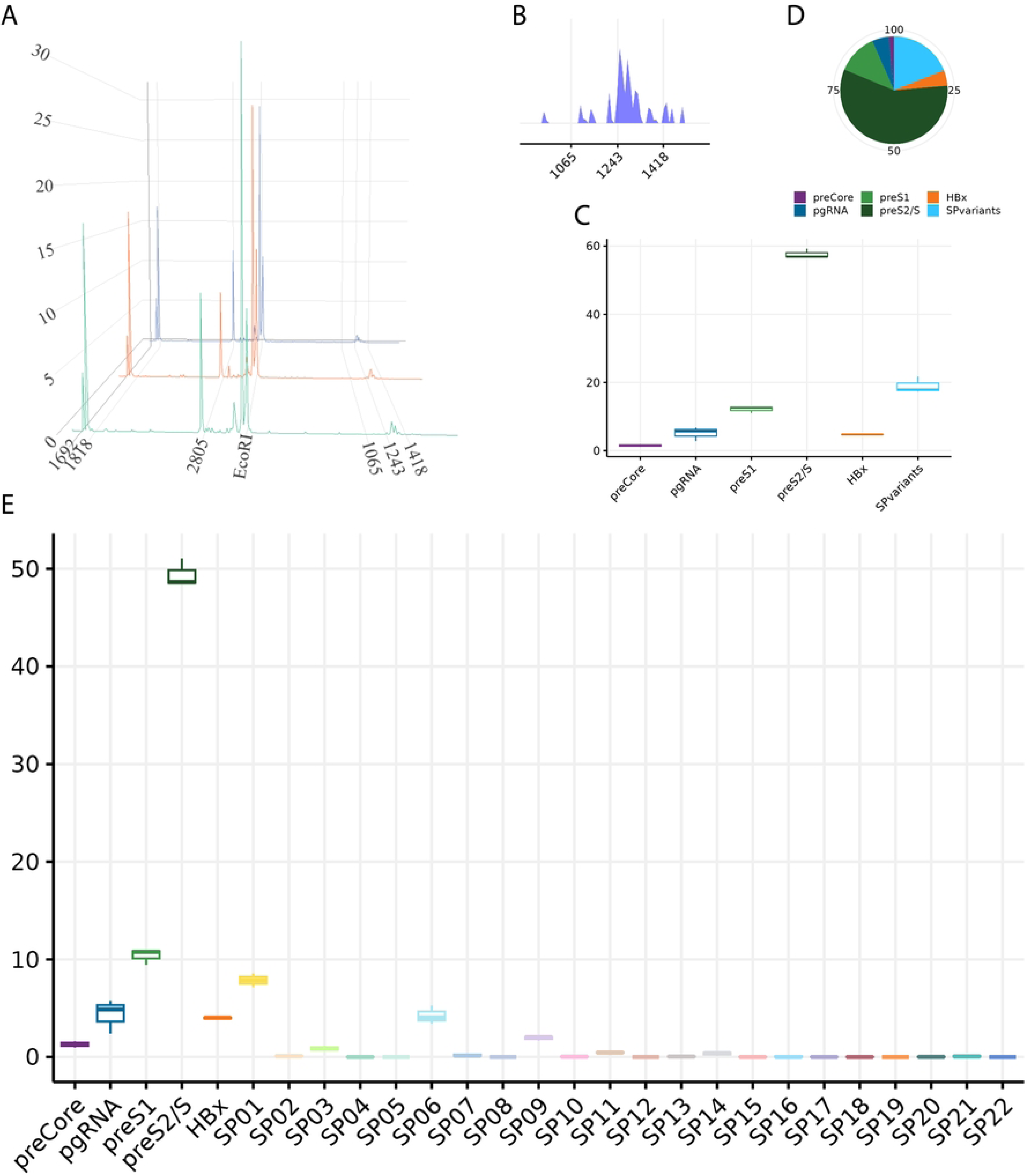
HBV RNA expression in HepG2-NTCP cells. HepG2-NTCP cells were infected with HBV for eight days. Total RNAs were extracted and subjected to HBV-specific 5’RACE-PCR followed by ONT sequencing. Sequencing data were analysed using Bolero. (A) Start position distributions of the three independent replicates (1, blue; 2, orange; 3, green). (B) Distribution of start positions of reads in a range of 1000 and 1500, that corresponds to putative L-, M- and S-HBx, in replicate 1. (C) Boxplot of reads proportion assigned to HBV RNAs based on putative TSS and splice junction identification. (D) Pie chart of read proportions assigned to HBV RNAs based on putative TSS and splice junction identification in replicate 1. (E) Boxplot of read proportions assigned to HBV RNAs based on putative TSS and splice junction identification.

We also took into account the TSS positions of HBx isoforms (Fig 10B), as their precise locations have remained uncertain in the field [16]. Although start position intervals for the three artificial HBx isoforms appeared clearly separated in simulated datasets (Fig 4D and S5 Fig), their discrimination in biological samples is less straightforward. As a result, we decided to adopt a single interval encompassing all HBx isoforms, spanning positions 1062 to 1519, rather than attempting to distinguish between HBx isoforms (Table 3). For each of the three independent replicates, reads were assigned to RNA species based on the TSS position ranges previously defined (S12 Fig). Subsequently, we calculated the mean percentage of reads assigned to a given TSS across the three replicates (S1 Table). These results provided evidence supporting the comparable levels of reproducibility as in simulated datasets, with the proportions for each RNA species remaining consistent across biological replicates. Analysis of the HBV transcriptome showed that the predominant RNA species, based on putative TSS identification, are preS2/S, accounting for 55% in mean of the total reads, then pgRNA (20% - including spliced variants), preS1 (15%), preCore (6%) and HBx (4%).

Next, splice junctions were detected in each read from the three independent replicates. On average, spliced variants represented 19% of the total number of reads (Fig 10C, D and S13 Fig). Upon closer inspection, spliced variants SP01, SP06, and SP09 were the most abundant ones. Nevertheless, almost all known spliced variants were detected at least once in the three independent replicates, with the exception of SP05, SP08, SP12, SP19 and SP22 (Fig 10E).

Finally, reads that cannot be categorized are placed in the “undefined” category. Apart from the 0.01% of reads that are removed to precisely identify spliced variants, this can be due to 1) a read’s start position falling outside the defined TSS ranges or 2) a splice junction or a combination of splice junctions that have not been reported yet. This approach thus has the potential advantage of being able to identify novel spliced variants.

## Discussion

Analysis of the HBV transcriptome remains challenging due to the particular structure of the viral genome. The organization of this circular DNA is highly condensed and all transcripts are, to various degrees, subsets of each other. Indeed, all HBV transcripts share the same 3’ end, and thus, the sequence of the shortest transcript, HBx, overlaps completely to the other transcripts. Moreover, preCore and pgRNA are longer than the viral DNA genome and are thus characterized by a 238-bp redundant region at both their 5’ and 3’ ends. Due to these constraints, HBV RNAs and spliced variants are indistinguishable by classical approaches, such as quantitative real time PCR or short-read transcriptome sequencing technologies. Indeed, in a large majority of cases, it is impossible to assign a short-read to a specific transcript. The exclusive discerning criterion for the identification of canonical HBV RNAs lies in the determination of the TSS position. In addition to the TSS, the presence of splice junctions serves as discernible feature for spliced variants.

In order to overcome these obstacles, long-read sequencing technologies have been employed to analyse and quantify the HBV transcriptome from both cell culture or patient serum samples [17,18].

Through ONT direct RNA and PCR-amplified cDNA sequencing, Vachon and co-workers detected HBV RNAs amidst total RNA extracted from chronic hepatitis B patients’ sera [18]. Direct RNA sequencing was unsuccessful due to low quantity of reads and PCR-amplified cDNA sequencing found minimal quantities of HBV RNAs, a median quantity of HBV reads of 0.08% of total reads. This approach offers a rapid assessment of HBV RNAs’ presence in patient serum. However, it remains unclear whether it is able to detect, distinct and identify every different known HBV RNA species and if it boasts sufficient sensitivity to detect extremely rare isoforms of spliced HBV RNA variants against the background of copious host RNAs.

According to an alternative approach proposed by Ng *et al*., the combination of an HBV RNA enrichment method based on biotinylated affinity probes with PacBio sequencing was developed [15]. Notably, their findings were consistent with ours, as the utilization of HBV-infected HepG2-NTCP cells enabled the identification of preS2 and HBs as the most prevalent viral RNAs and the detection of common spliced variants, including SP01 (S2 Table). However, significant discrepancies were observed in the quantification of preCore and HBx RNA species. Specifically, Ng and colleagues detected 13% and 0.2% on average for preCore and HBx, respectively, whereas our study detected 1.3% and 4.7%, respectively, for these two RNA species. Furthermore, regarding spliced variants, Ng and colleagues detected 9% on average, while more than 20% were detected in our work. Nevertheless, the most represented spliced variants in both analyses were SP01, SP06, and SP09. These quantitative differences could be attributed to several factors. Firstly, different HBV RNA enrichment methods were employed in the two studies. While Ng *et al.*’s method captures HBV RNAs via affinity from total RNA extractions, our method specifically isolates and amplifies mature, capped, and full-length HBV RNAs through the 5’RACE method [5]. Additionally, there may be variations in efficiency and selectivity between these two methods, leading to differing sequencing libraries. Secondly, although both experiments used HepG2-NTCP cells, slightly different culture methods were employed in each study. Moreover, distinct strains of HBV were used for infection in the two works, potentially resulting in varying levels of expression of their RNAs and spliced variants. Regarding preCore RNA, Ng *et al*., distinguished between long and short preCore forms. There might be a difference in assignment of these RNAs between their method and ours. Similarly, it should be noted that spliced variant SP12 was identified as pSP12 (partial-SP12) by Ng and colleagues, representing a truncated form of this variant. Given that the method associated with Bolero excludes truncated RNAs, this could explain why spliced variant SP12 was not detected in our investigation.

In comparison to the approaches outlined by Ng *et al.* and Vachon *et al.*, Bolero has been conceptualized and evaluated using simulated sequencing data that incorporates all reported HBV RNAs. The prediction suggests that Bolero can accurately detect and precisely assign these RNA species. Moreover, our method harnesses ONT, which generally provides lower costs relative to PacBio sequencing. Furthermore, potential inaccuracies arising due to sequencing errors can be significantly reduced owing to the recent development of a new chemistry for ONT sequencing technology [17,18].

In addition, the 5’RACE method developed by Stadelmayer *et al.* offers the capability to exclusively amplify RNAs transcribed from covalently closed circular DNA (cccDNA) within the context of hepatitis B virus (HBV) research. Through strategic placement of the gene-specific primer (GSP1; positions 1790-1809) downstream of the direct repeat 2 (DR2) integration site (positions 1591-1602), the original design ensures selective capturing and amplification of cccDNA-generated RNAs while excluding those derived from potentially integrated HBV sequences within the host genome. However, recognizing the importance of understanding the contribution of both sources, the 5’RACE approach can be further tailored to encompass all HBV RNAs stemming from either cccDNA or integrated sequences. An illustrative example showcases the utilization of an alternative gene-specific primer (GSP2; positions 1566-1586) situated upstream of the DR2 integration site for performing the initial reverse transcription PCR (RT-PCR) step and subsequent 5’RACE amplification steps. This capability to distinguish cccDNA-generated RNAs from those produced from integrated sequences represents a significant strength of Bolero.

Finally, Bolero not only enables the quantification of HBV RNA species, but also stores information of putative TSS associated with particular splice junctions for each read, allowing the potential identification of novel spliced variants that haven’t yet been described in the literature. Thus, our work shows that Bolero is highly efficient for the discrimination and quantification of individual HBV mRNAs, as well as a relevant tool to better understand the complex molecular mechanisms regulating the HBV transcriptome.

## Material and methods

### Workflow implementation

The Bolero workflow is implemented as a set of open-source and free to use software (with exception of the ONT software guppy, which is facultative [21]), Python and R modules, fully automated using Nextflow management system [22]. All software and scripts, including the dependencies are automatically managed by Nextflow, taking advantage of the containerization using Docker technology. Docker images are available at https://hub.docker.com/, and can be automatically converted in singularity or charliecloud images with Nextflow. Containerization ensures reproducibility and facilitate the usage of the workflow.

### Basecalling, preprocessing and trimming for Nanopore adapters

The workflow scheme is linear (Fig 2), handling Fast5 files from Oxford Nanopore software, such as MinKnow Software, or basecalled reads Fastq files as input. Users have the possibility to basecall the raw sequencing data using guppy in cpu or gpu mode. A filtering of low-quality reads is setup during the basecalling step to exclude reads with a Qvalue ≤ 7. The ONT sequencing kit and flowcell parameters are required to set demultiplexing step. If the Fastq files are provided, the basecalling step is skipped and the filtering of reads for their quality is under the responsibility to the user. Default parameters are set to skip the basecalling and demultiplexing steps but are tuneable using Nextflow command line passing options. Sequencing adapters are trimmed using porechop (v.0.2.4) [23].

### Read filtering

Trimmed reads containing firstly the 5’RACE adapter sequence and then the GSP used in the 5’RACE amplification step are filtered in using Seqkit software (v.2.4.0) [8]. 5’RACE and GSP sequences are provided by the user. To cope with ONT sequencing error rate, 10% of mismatches are allowed in 5’RACE adapter and GSP sequences detection using the *--max-mismatch* option of *seqkit grep*. Then, 5’RACE adapter is trimmed using cutadapt (v.2.8) [24].

### Mapping of reads

Filtered and trimmed reads were aligned to the user-provided reference genome using minimap2 (v.2.17) [10,11]. Here, the 3421-nucleotide long preCore RNA sequence of the HBV genotype D (ayw strain, NC_003977.2), from the TSS (i.e., 122 nucleotides upstream the pgRNA start codon) to the PAS were used as reference sequence. The dedicated preset *spliced long reads* mode is set with the *-ax splice* option. In order to quantify the expression of the different HBV RNA species, no secondary alignments are allowed, using the minimap2 option *--secondary=no*. Concerning splicing events, the detection of non-canonical splicing sites, no attempt to match GT-AG, is set with the option *-u n* and *--splice-flank=no*. Finally, the output =/X CIGAR operators for sequence *match/mismatch* option is set with the *-eqx* parameter. Mapped reads are stored, binarized, sorted and indexed in BAM and BAI files using samtools (v.1.17) [11–13].

### Simulation of reads

An HBV 5’RACE sequencing by ONT was simulated by the PBSIM3 software (v.3.0.0) [25]. Using the Hidden Markov error model (errhmm) and the transcriptome mode provided by the tool, several datasets of individual known HBV RNA species, including SP01 to SP22 spliced variants, are simulated in balanced reads number per dataset. Distinct sequencing depths are simulated to evaluate the variability of reads and the impact of ONT error rate. The range of depth is dispersed between 10^4^ to 10^6^ reads per RNA species (data not shown). Optimizations were established using datasets of 2.5×10^5^ simulated reads per HBV RNA species and spliced variants to obtain a comparable synthetic dataset, in terms of quantity of preprocessed reads obtained by sequencing cell lines. Finally, a complete transcriptome, is obtained by the combination of all individual datasets. A triplicate of synthetic datasets has been taken into account to determine TSS and splicing site position ranges. The simulation produces artefactual reads shorter than expected, hence, in addition to workflow filtering steps, synthetic reads shorter than 390 bp are filtered-out.

### Biological sample preparation, 5’RACE and sequencing

HepG2-NTCP cells were cultured in DMEM medium (Life Technologies, Carlsbad, CA, USA) supplemented with 5% fetal calf serum (FCS, Cytiva, Marlborough, MA, USA). Two days prior to infection, 2.5% DMSO was added. Cells were then infected with 250 viral genome equivalents/cell in infection medium (DMEM supplemented with 5% FCS and 2.5% DMSO) in presence 4% polyethylene glycol (PEG 8000). One- or four-days post-infection, cells were washed three times with 1X PBS and cultured during four extra days in infection medium. RNAs were extracted using RNAzol (MRC) according to the manufacturer’s instructions and subjected to 5’RACE-PCR amplification as previously described [5]. 5’RACE-PCR library was then prepared using the ligation sequencing gDNA-PCR barcoding kits (SQK-LSK109 with EXP-PBC001) and sequenced on a MinION device (ONT^®^). The device was driven by MinKnow (ONT^®^).

## Author contributions

Conceptualization: XG, DB, BT

Data Curation: XG, AR

Formal Analysis: XG, AR

Funding Acquisition: BT, FZ, ML

Investigation: GG, AP, DB

Methodology: XG

Project Administration: BT

Software: XG, AR

Supervision: BT, CB, FZ

Visualization: XG, GG

Writing – Original Draft Preparation: XG, GG, AARS, BT

Writing – Review & Editing: All authors

## Data and code availability

Complete workflow is open source and free-to-use, under the AGPLv3.0. Code and documentation are available on GitLab (https://gitbio.ens-lyon.fr/xgrand/bolero).

## Acknowledgements

The authors would like to thank the CBP and the PSMN (https://www.ens-lyon.fr/PSMN/doku.php?id=en:accueil) to provide high-performance computational facilities, HubBioinfo https://www.ens-lyon.fr/LBMC/pole-bioinformatique?set_language=en&cl=en, and particularly Laurent Modolo, Nicolas Fontrodona and Hélène Polvèche from Ecole Normale Supérieure de Lyon for their informatics and bioinformatics trainings, expertise sharing and advices.

## Supplemental information

**S1 Fig. Distribution of start positions for each individual HBV RNA species in synthetic datasets (replicate 1).** Positions are EcoRI based on the HBV genome strain ayw, genotype D.

**S2 Fig. Distribution of start positions for each individual HBV RNA species in synthetic datasets (replicate 2).** Positions are EcoRI based on the HBV genome strain ayw, genotype D.

**S3 Fig. Distribution of start positions for each individual HBV RNA species in synthetic datasets (replicate 3).** Positions are EcoRI based on the HBV genome strain ayw, genotype D.

**S4 Fig. Distribution of start positions for combined preCore and pgRNA in synthetic datasets (replicates 2 and 3).** Positions are EcoRI based on the HBV genome strain ayw, genotype D. Purple and blue dashed lines represent minimum and maximum start positions of preCore and pgRNA synthetic reads respectively.

**S5 Fig. Distribution of start positions for combined L-, M- and S-HBx isoforms in synthetic datasets (replicates 2 and 3).** Positions are EcoRI based on the HBV genome strain ayw, genotype D. Orange, red and dark red dashed lines represent minimum and maximum start positions of L-, M- and S-HBx isoforms synthetic reads respectively.

**S6 Fig. Distribution of start positions for individual preS2 and HBs in synthetic datasets (replicates 2 and 3).** Positions are EcoRI based on the HBV genome strain ayw, genotype D. Light and dark green dashed lines represent minimum and maximum start positions of preS2 and HBs synthetic reads respectively.

**S7 Fig. Distribution of start positions of complete synthetic datasets (replicates 2 and 3).**Positions are EcoRI based on the HBV genome strain ayw, genotype D.

**S8 Fig. Relative and absolute read quantities according to putative TSS groups in synthetic datasets (replicates 2 and 3).** Pie chart of assigned reads relative quantities to putative TSS groups. #reads represents the total number of assigned reads. Comparison of absolute quantities of reads assigned to a putative TSS by Bolero, based on start positions and preprocessed simulated ones as input of quantification step.

**S9 Fig. Density of reads expressed as a function of donor or acceptor site positions of splice junctions.** J2450_490 junction donor (red) and acceptor (blue) sites position distributions respectively, and position distributions of donor sites of J2450_490 and J2474_283 junctions. Plain vertical lines are minimum and maximum positions, dashed vertical lines are selected borders, defining interval containing 99.9% of donor or acceptor site positions.

**S10 Fig. Comparison of absolute read quantities assigned by Bolero and preprocessed simulated reads as input of the quantification step in synthetic datasets (replicates 2 and 3).** Quantification takes into account putative TSS of reads and the presence of splice junctions.

**S11 Fig. Relative expression of HBV RNA species in synthetic datasets (replicates 2 and 3).** Bar chart of relative quantification of known HBV RNA species, and pie chart of relative quantification of known canonical RNA species and spliced variants group.

**S12 Fig. Relative read quantities according to putative TSS groups in HBV-infected HepG2-NTCP cells.** HepG2-NTCP cells were infected with HBV for eight days. Total RNAs were extracted and subjected to an HBV-specific 5’RACE-PCR followed by ONT sequencing. Sequencing data were analysed using Bolero. Pie charts represent assigned reads relative quantities to putative TSS groups of the three independent replicates.

**S13 Fig. Relative read quantities in HepG2-NTCP infected cells.** HepG2-NTCP cells were infected with HBV for eight days. Total RNAs were extracted and subjected to an HBV-specific 5’RACE-PCR followed by ONT sequencing. Sequencing data were analysed using Bolero. Pie charts of read proportions assigned to HBV RNAs based on putative TSS and splice junction identification in independent replicates 2 and 3.

**S1 Table. Mean percentage of reads assigned by Bolero to a putative TSS across the three replicates of HBV infected HepG2-NTCP cells.**

S2 Table. Comparison of relative quantification of HBV RNA species in HepG2-NTCP cells obtained with Bolero, compared to those reported by Ng *et al.* 2023. Canonical HBV RNAs (preCore, pgRNA, preS1, preS2/S and HBx) quantification according to putative TSS from Ng *et al.*, compared to the ones obtained by Bolero. Proportion of spliced variants obtained by Ng *et al.,* compared to those obtained by Bolero, taking into account putative TSS and splice junctions.

## References

1. Polaris Observatory Collaborators. Global prevalence, treatment, and prevention of hepatitis B virus infection in 2016: a modelling study. Lancet Gastroenterol Hepatol. 2018;3: 383–403. doi:10.1016/S2468-1253(18)30056-6

2. Martinez MG, Boyd A, Combe E, Testoni B, Zoulim F. Covalently closed circular DNA: The ultimate therapeutic target for curing HBV infections. Journal of Hepatology. 2021;75: 706–717. doi:10.1016/j.jhep.2021.05.013

3. Testoni B, Scholtès C, Plissonnier M-L, Paturel A, Berby F, Facchetti F, et al. Quantification of circulating HBV RNA expressed from intrahepatic cccDNA in untreated and NUC treated patients with chronic hepatitis B. Gut. 2023 [cited 17 Feb 2024]. doi:10.1136/gutjnl-2023-330644

4. Altinel K, Hashimoto K, Wei Y, Neuveut C, Gupta I, Suzuki AM, et al. Single-Nucleotide Resolution Mapping of Hepatitis B Virus Promoters in Infected Human Livers and Hepatocellular Carcinoma. J Virol. 2016;90: 10811–10822. doi:10.1128/JVI.01625-16

5. Stadelmayer B, Diederichs A, Chapus F, Rivoire M, Neveu G, Alam A, et al. Full-length 5’RACE identifies all major HBV transcripts in HBV-infected hepatocytes and patient serum. Journal of Hepatology. 2020;73: 40–51. doi:10.1016/j.jhep.2020.01.028

6. Kremsdorf D, Lekbaby B, Bablon P, Sotty J, Augustin J, Schnuriger A, et al. Alternative splicing of viral transcripts: the dark side of HBV. Gut. 2021;70: 2373–2382. doi:10.1136/gutjnl-2021-324554

7. Wang Y, Zhao Y, Bollas A, Wang Y, Au KF. Nanopore sequencing technology, bioinformatics and applications. Nat Biotechnol. 2021;39: 1348–1365. doi:10.1038/s41587-021-01108-x

8. Shen W, Le S, Li Y, Hu F. SeqKit: A Cross-Platform and Ultrafast Toolkit for FASTA/Q File Manipulation. PLoS One. 2016;11: e0163962. doi:10.1371/journal.pone.0163962

9. Bichko V, Pushko P, Dreilina D, Pumpen P, Gren E. Subtype ayw variant of hepatitis B virus. DNA primary structure analysis. FEBS Lett. 1985;185: 208–212. doi:10.1016/0014-5793(85)80771-7

10. Li H. Minimap2: pairwise alignment for nucleotide sequences. Bioinformatics. 2018;34: 3094–3100. doi:10.1093/bioinformatics/bty191

11. Li H. New strategies to improve minimap2 alignment accuracy. Bioinformatics. 2021;37: 4572–4574. doi:10.1093/bioinformatics/btab705

12. Li H, Handsaker B, Wysoker A, Fennell T, Ruan J, Homer N, et al. The Sequence Alignment/Map format and SAMtools. Bioinformatics. 2009;25: 2078–2079. doi:10.1093/bioinformatics/btp352

13. Danecek P, Bonfield JK, Liddle J, Marshall J, Ohan V, Pollard MO, et al. Twelve years of SAMtools and BCFtools. GigaScience. 2021;10: giab008. doi:10.1093/gigascience/giab008

14. Delahaye C, Nicolas J. Sequencing DNA with nanopores: Troubles and biases. PLoS One. 2021;16: e0257521. doi:10.1371/journal.pone.0257521

15. You Y, Clark MB, Shim H. NanoSplicer: accurate identification of splice junctions using Oxford Nanopore sequencing. Bioinformatics. 2022;38: 3741–3748. doi:10.1093/bioinformatics/btac359

16. Peng B, Jing Z, Zhou Z, Sun Y, Guo G, Tan Z, et al. Nonproductive Hepatitis B Virus Covalently Closed Circular DNA Generates HBx-Related Transcripts from the HBx/Enhancer I Region and Acquires Reactivation by Superinfection in Single Cells. J Virol. 2023;97: e0171722. doi:10.1128/jvi.01717-22

17. Ng E, Dobrica M-O, Harris JM, Wu Y, Tsukuda S, Wing PAC, et al. An enrichment protocol and analysis pipeline for long read sequencing of the hepatitis B virus transcriptome. J Gen Virol. 2023;104: 001856. doi:10.1099/jgv.0.001856

18. Vachon A, Seo GE, Patel NH, Coffin CS, Marinier E, Eyras E, et al. Hepatitis B virus serum RNA transcript isoform composition and proportion in chronic hepatitis B patients by nanopore long-read sequencing. Front Microbiol. 2023;14: 1233178. doi:10.3389/fmicb.2023.1233178

19. Zhang T, Li H, Ma S, Cao J, Liao H, Huang Q, et al. The newest Oxford Nanopore R10.4.1 full-length 16S rRNA sequencing enables the accurate resolution of species-level microbial community profiling. Applied and Environmental Microbiology. 2023;89: e00605-23. doi:10.1128/aem.00605-23

20. Lerminiaux N, Fakharuddin K, Mulvey MR, Mataseje L. Do we still need Illumina sequencing data? Evaluating Oxford Nanopore Technologies R10.4.1 flow cells and the Rapid v14 library prep kit for Gram negative bacteria whole genome assemblies. Can J Microbiol. 2024; cjm-2023-0175. doi:10.1139/cjm-2023-0175

21. Wick RR, Judd LM, Holt KE. Performance of neural network basecalling tools for Oxford Nanopore sequencing. Genome Biol. 2019;20: 129. doi:10.1186/s13059-019-1727-y

22. Di Tommaso P, Chatzou M, Floden EW, Barja PP, Palumbo E, Notredame C. Nextflow enables reproducible computational workflows. Nat Biotechnol. 2017;35: 316–319. doi:10.1038/nbt.3820

23. Wick RR, Judd LM, Gorrie CL, Holt KE. Completing bacterial genome assemblies with multiplex MinION sequencing. Microb Genom. 2017;3: e000132. doi:10.1099/mgen.0.000132

24. Martin M. Cutadapt removes adapter sequences from high-throughput sequencing reads. EMBnet.journal. 2011;17: 10–12. doi:10.14806/ej.17.1.200

25. Ono Y, Hamada M, Asai K. PBSIM3: a simulator for all types of PacBio and ONT long reads. NAR Genom Bioinform. 2022;4: lqac092. doi:10.1093/nargab/lqac092

